# Controllable drug release of pH-sensitive liposomes encapsulating artificial cytosol system

**DOI:** 10.1101/2021.05.24.445400

**Authors:** Wei Zong, Xiaotong Shao, Yunhe Chai, Xiuwen Wang, Shuang Han, Hongtao Chu, Chuntao Zhu, Xunan Zhang

## Abstract

The fabrication of cell model containing artificial cytosol is challenging. Herein we constructed an artificial cytosol contained cell model by electroformation method. Agarose was selected as the main component of the artificial cytosol. Sucrose was added into agarose to regulate the sol viscosity and phase transition temperature (*T*_m_). The viscosity of the sol with the mass ratio (agarose-sucrose) 1:9 was closest to the natural cytosolic. DSPC/20 mol% Chol was used to form large unilamellar vesicle (LUV) as cell model compartment. The rhodamine release experiment confirmed that the release of rhodamine from LUVs containing artificial cytosol took more time than that from LUVs containing pure water. The unique release profile makes agarose-sucrose@LUVs suitable as a drug carrier. Doxorubicin (DOX) is loaded in the agarose-sucrose@LUVs, and their half maximum inhibition concentration on HeLa cells is 0.015 μmol L^−1^, which means 31.7 times increase in inhibition efficiency over free DOX.

## 1. Introduction

Cell is the basic structural and functional unit of all known living organisms.[1] The cell is one of the most complex and exciting systems in nature that scientists are still trying to fully understand. However, along with the rapid development of cell biology, many issues have arisen, such as the inherent complexity and frangibility of biological cells, that is to say, easy loss of activity or death in vitro. To overcome these issues while still mimicking biological cells, artificial cells are built[2], which are expected to be more easily controlled and more robust than natural cells. These artificial cells can have applications in many fields from medicine to environment, and may be useful in constructing the theory of the origin of life.[3] Vesicular assemblies of phospholipids are perfect drug carriers because the inherent advantages such as biocompatibility, low toxicity, high loading capacity, and controllable release kinetics.[4, 5] Large unilamellar vesicle (LUV) is more suitable as a drug carrier than gaint unilamellar vesicle (GUV). Phospholipid vesicles are also called liposomes. Liposomes have demonstrated a wide range of clinical applications ranging from therapeutic, diagnostic to theranostic applications.[5] The liposomes loaded with anticancer drugs can accumulate inside the target tumor tissue via passive targeting. This phenomenon is highly manifested in solid tumors via a process known as enhanced permeation and retention (EPR) effect.[6]

Regarding cell biomimicry, in addition to mimic the cell membrane and organelles, cytoplasm simulate is also be studied in progress. Cytoplasm plays an important role in cellular activity and regulation[7]. Cytoplasm mainly consists of cytosol and organelles. Cytosol is highly crowded liquid or gel-like phase because it dissolves high concentrations of molecules. Therefore, scientists construct various crowding system that mimic the cytosol environment to study the biochemical reactions that take place in real cells. Saccharides with low molecular weight have been used to create an effective crowding microenvironment for gene expression in cell-free protein synthesis (CFPS). At a low concentration range of saccharides, the mRNA and protein amounts will be on the rise with the concentration increment of saccharide. When the concentrations exceed a certain value, the mRNA and protein amounts decrease. This provides a new sight that biochemical reactions would be affected by substances with low molecular weight in the cellular physical environment.[8] Huck et al. produced an artificial cell-like environment consisting of condense Escherichia coli lysate for gene expression. Compared with noncrowded conditions, the binding constant of T7 RNA polymerase to DNA and the transcription rate constant were significantly enhanced under crowded conditions. These results enable us to further understand the effects of crowding on key cellular processes such as transcription and translation in living system.[9] Nanoclay is a novel nanomaterial, which was applied to create a locally crowded environment to explore its effects on protein expression in the cell-free system. A higher concentration in the cell-free system was formed due to the adsorption effect between the plasmid and nanoclay surface. Under such crowded environment, protein expression was successfully achieved in the cell-free system and the process of translation and transcription were improved simultaneously. This work provides us with a new idea that nanomaterials could be applied to mimic cellular microenvironment.[10] Vesicles with a gelly or gelified cavity, also named “hydrosomes”, have been investigated for approximately 20 years. There are many interesting experimental results arising when mimicking a viscoelastic cytosol, such as a slower diffusion rate of molecules from inside of membrane toward the outside environment. Due to the mechanical stress, artificial cell stability and shape integrity can get a better protection. There are many protocols for constructing artificial cells containing cytosol, but one is often used, consists in encapsulating a poly(N-isopropylacrylamide) (PNIPAAm)-based solution, which will in situ photopolymerize into covalently cross-linked gels under UV irradiation.[11–18] In another work, a poly(ethylenedioxythiophene) (PEDOT)/PSS polyelectrolyte mixture previously microinjected in liposomes was cross-linked by electrostatic complexation via Ca^2+^ influx.[19] In nature, the cytoskeleton is composed of the protein filaments actin, microtubules and intermediate filaments, formed by nucleation elongation processes. The growth of the protein filaments of this cytoskeleton is driven by noncovalent interactions, making them very dynamic with constant association/dissociation processes.[20] So noncovalent or physical gels constitute in our opinion a better alternative.

The choice of materials for building artificial cell is crucial. The phospholipid membrane can actually simulate the structure of nature cell membrane and the transfer relationship of micro reaction in artificial cells.[21] There have been many studies using liposomes to construct artificial cells or organelles, such as artificial eukaryotes[22, 23], artificial mitochondria[24], artificial chloroplasts[25, 26], and so on. Herein, we demonstrated the formation of agarose-sucrose@LUVs. Using agarose and sucrose gel to simulate the viscosity of artificial cytosol and liposome to mimic cell compartment. The molecule diffusion rate and morphological stability of agarose-sucrose@LUVs were investigated. Compared to the exponential drug release of common pH-sensitive LUVs, the novel agarose-sucrose@LUVs possess a unique slowly release profile. Furthermore, the new system exhibits better inhibition efficiency than common pH-sensitive LUVs, because of longer release time provided by agarose-sucrose release, and higher uptake rate of doxorubicin (DOX). The new system is considerably stable. From all above, the agarose-sucrose@LUVs system is quite competent as drug delivery system.

## 2. Materials and Methods

### 2.1 Materials

1,2-dimyristoyl-sn-glycero-3-phosphocholine (DMPC), 1,2-dipalmitoyl-sn-glycero-3-phosphocholine (DPPC), 1,2-Dioctadecanoyl-sn-glycero-3-phophocholine (DSPC), cholesterol (Chol) were purchased from Avanti Polar Lipids (USA). Fluorescence-labelled 1,2-dioleoyl-sn-glycero-3-phosphoethanolamine-N-(7-nitro-2-1,3-benzoxadiazol-4-yl) (NBD-PE) was obtained from Molecular Probes (Eugene, Oregon, US). Sucrose was purchased from XiLong Chemical Co., Ltd. (China) Glass slides coated with indium tin oxide (ITO, sheet resistance ≈ 8 to 12 Ω, thickness ≈ 160 nm) were purchased from Hangzhou Yuhong technology Co. Ltd (China). Agarose, chloroform and absolute ethanol were purchased from Sigma (China). Millipore Milli-Q water with a resistivity of 18.0 MΩ cm was used for solution preparation.

### 2.2 Preparation of agarose and sucrose sol

Agarose and sucrose solid powders were weighed. The total mass of solid powder was controlled at 60 mg and the volume of liquid water was 1 mL. Various agarose sols concentrations were prepared, which is 0.056% w/w (pure agarose), 0.028% w/w (1:1, agarose/sucrose, w/w), 0.019% w/w (1:2, agarose/sucrose, w/w), 0.014% w/w (1:3, agarose/sucrose, w/w), 0.011% w/w (1:4, agarose/sucrose, w/w), 0.009% w/w (1:5, agarose/sucrose, w/w), 0.007% w/w (1:7, agarose/sucrose, w/w), 0.005% w/w (1:9, agarose/sucrose, w/w), 0.004% w/w (1:14, agarose/sucrose, w/w), 0.003% w/w (1:19, agarose/sucrose, w/w). They were first homogenized at 100℃ during at least 1 h. The sols then slowly cooled down. The critical temperature at which the sol converts to gel is studied and an optimal concentration was selected for the artificial cytosol.

### 2.3 Encapsulation agarose-sucrose gel in GUVs

Electroformation is an especially attractive method to generate GUVs since it provides higher yield with short time period. ITO-coated glass coverslips (25 mm × 45 mm) were cleaned in ethanol and water each for 15 minutes by sonication and then dried by N_2_. Lipid of DSPC/20 mol% Chol including additional fluorescence lipid NBD-PE was dissolved in chloroform to make a solution concentration of 5 mg mL^−1^. The lipid solution was deposited on ITO electrode surface using a needle to spread carefully back and forth 6 times, followed by drying under vacuum for 2 h. The coverslips were separated by a rectangular polytetrafluoroethylene (PTFE) spacer with a length, width and height of 35 mm, 25 mm and 2 mm, respectively. The AC-electric field was applied for to generate GUVs containing agarose-sucrose mixed sol.

### 2.4 Encapsulation agarose-sucrose gel in LUVs

Artificial cytosol-LUVs were produced by mini-extruder. The lipid stocking solution was dried in a test tube and then dissolved in DOX loaded artificial cytosol, followed by vortexing for 3 min, preheating at 55 °C. The solution then went through the polycarbonate membranes with pores of 200 nm diameter for 11 times back and forth with syringes. The obtained LUVs are about 160 nm. The LUV suspension was also purified with the same method as GUVs.

### 2.5 Estimation of stability

Stability of agarose-sucrose@LUVs and DOX loaded agarose-sucrose@LUVs were evaluated by dynamic light scattering (DLS), polymer dispersity index (PDI) and ζ-potential. DLS was tested using a material refringence of 1.590 and a dispersant index of 1.33.

### 2.6 In vitro controlled release of DOX-agarose-sucrose@LUVs

The in vitro release of DOX-agarose-sucrose@LUVs was performed by a dialysis method. The DOX-agarose-sucrose@LUV suspension (0.5 mL) was added into a dialysis membrane (molecular weight cut-off 8–14 kDa, MD 44, Solarbio, China) that was subsequently placed into 20 mL buffer solution (sodium hydrogen phosphate-citric acid, pH = 5 and pH = 7.4 respectively) under magnetic stirring at 37 °C. At desirable time intervals, 2 mL of supernatant were taken to measure fluorescent intensity and then returned back to mother solution. The release percentage at each time interval was obtained by the amount of DOX released at that time over the total amount of DOX inside the LUVs.

### 2.7 Cell culture and viability

The cells were grown in a RPMI-1640 (GIBCO) medium with 10% (v/v) fetal bovine serum (FBS). The cells were maintained in an incubator at 37 °C in an atmosphere of 5% CO_2_. The cells were seeded in 96-well plates with a density of 6000 viable cells per well and incubated for 24 h for cell attachment. The antitumor activity of the compounds was tested by standard MTT experiments. Various concentration of DOX solutions were prepared. After 24 h, 20 μL MTT solution at a concentration of 5 mg mL^−1^ dissolved in phosphate buffer solution (PBS, pH 7.4) was added to each well. After another 4 h incubation, the medium was removed and 100 μL dimethylsulfoxide (DMSO) was added into each well. The intensity of the absorbance was measured using an automatic microplate reader at a wavelength of 495 nm. The results were expressed as mean values ± standard deviation of 3 measurements.

### 2.8 Uptake of DOX-agarose-sucrose@LUVs by HeLa cells and their localization in cells

To evaluate the uptake of DOX-ATPS-LUVs by HeLa cells, lipid was labelled with NBD-PE. Cells were seeded in a 24-well plate in RPMI-1640 medium overnight. Fluorescence-labelled liposomes were suspended over a bath sonicator for 3 min, and 0.5 mL solution was added to the cultured cells. After a certain period of time, the cells were fixed with 4% paraformaldehyde for 10 min and washed three times with PBS, then stained with DAPI (100 ng mL^−1^) for 3 min. After washing with PBS, the coverslips were mounted immediately and cells were directly visualized by a fluorescence microscope.

## 3. Results and discussion

### 3.1 Modulation of the critical temperature and sol viscosity

The critical temperature of agarose (from sol to gel) can be easily tuned from 40°C (temperature transition in pure water) to 20°C by adjusting the sucrose concentration within the sol. After homogenized heating at 100°C for 1 h, the sol then slowly cooled down to 37°C. The variation of sol states is as shown in Fig. 1a. The sample presents a creamy white jelly state when the mass ratio of agarose/sucrose were 1:0, 1:1, 1:2. When the mass ratio of agarose/sucrose was 1:3 the sample presents semifluid state. The sample showed the flow state when the mass ratio of agarose/sucrose were 1:4, 1:5, 1:7, 1:9, 1:14, 1:19, respectively. The inverted bottles can easily present the colloidal state as shown in the inset of Fig. 1a. The gel was all curdled at the bottom of the bottle when the mass ratio of agarose/sucrose was 1:2. One part of the gel solidified at the bottom of the bottle, and the other part flowed to the mouth of the bottle when the mass ratio of agarose/sucrose was 1:3. The sol can all flow to the mouth of the bottle when the mass ratio of agarose/sucrose was 1:4, also 1:5, 1:7, 1:9, 1:14 and 1:19, respectively. The transition temperature of agarose and sucrose mixed sol gradually decreased along with increasing of the concentration sucrose as shown in Fig. 1b. Their critical temperature from sol to gel were 39.9±0.85℃, 38.6±0.48℃, 37.9±0.26°C, 37.3±0.48°C, 36.5±0.50°C, 34.2±0.29°C, 32.5±0.55°C, 29.7±0.95°C, 27.8±0.55°C and 24.1±0.34°C for the mass ratio of agarose and sucrose were 1:0, 1:1, 1:2, 1:3, 1:4, 1:5, 1:7, 1:9, 1:14 and 1:19, respectively.

**Fig. 1.**
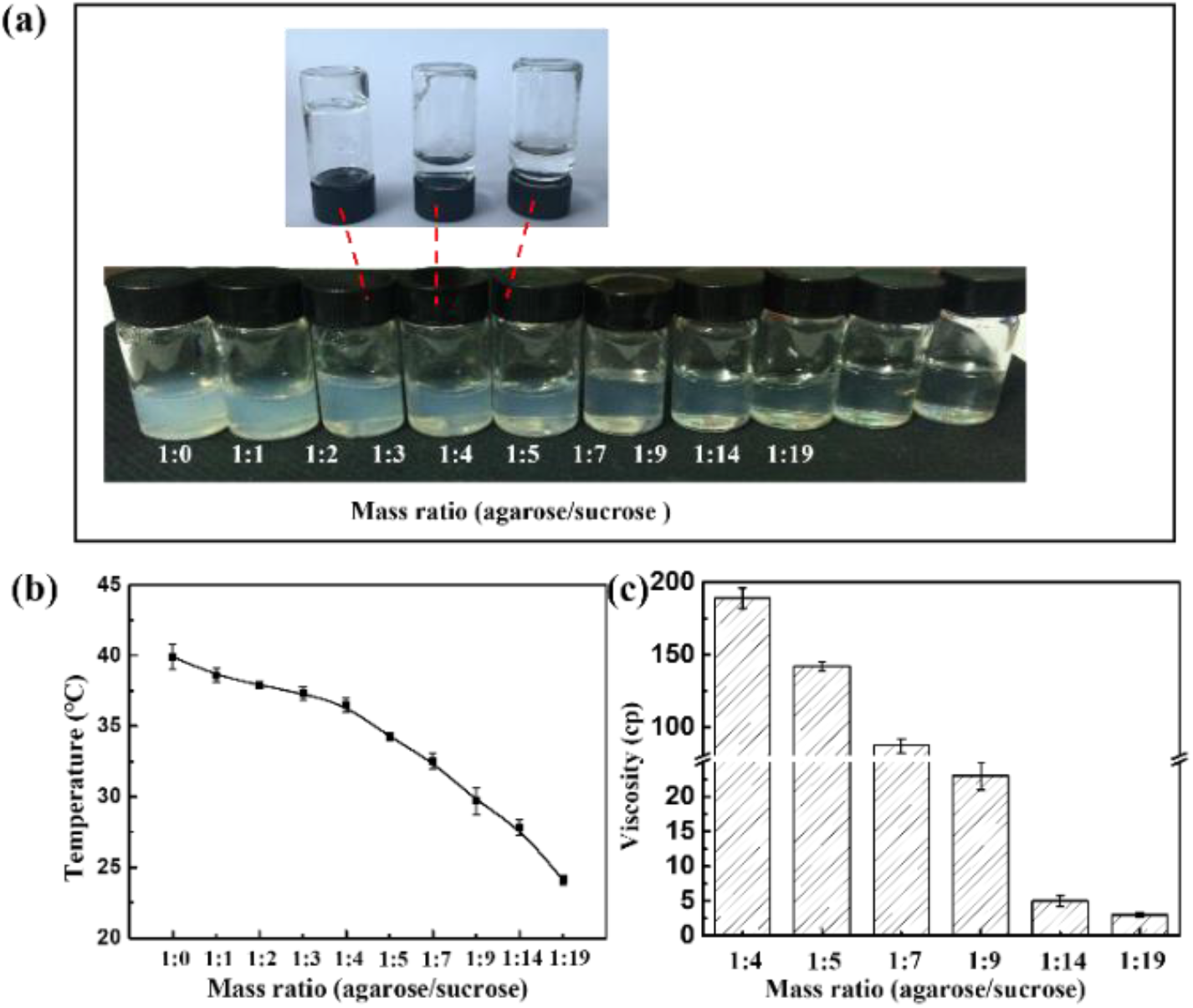
(a) The images of agarose-sucrose mixed sol cooled to 37°C. (b) Critical temperature of agarose-sucrose (from sol to gel). (c) The viscosity of agarose-sucrose mixed sol at 37°C.

The viscosity of sample that the mass ratio (agarose/sucrose) at 1:4, 1:5, 1:7, 1:9, 1:14, and 1:19 were measured by viscometer at 37°C, as shown in Fig. 1c. The viscosity of agarose-sucrose mixed sol gradually decreased along with increasing of the concentration of sucrose. The viscosity was 189±7 cp, 142±3 cp, 87±5 cp, 23±2 cp, 5±0.8 cp, and 3±0.3 cp for the mass ratio of agarose and sucrose were 1:4, 1:5, 1:7, 1:9, 1:14 and 1:19, respectively. Typical viscosity of cytosol ranges from 5 to 30 cp, and it has been postulated that they play an important role in cell communication processes.[27–29] The viscosity of the mass ratio **(agarose/sucrose)**at 1:9 was closest to the nature cytosolic viscosity. Therefore, agarose/sucrose mixed sol which the mass ratio was 1:9 was chose as artificial cytosol in this paper.

### 3.2 Preparation of the agarose-sucrose@GUVs

DMPC (*T*_m_=24°C), DPPC (*T*_m_=41°C) and DSPC (*T*_m_=55°C) GUVs in artificial cytosol were produced as mentioned above, respectively. Fig. S1 showed typical fluorescence images of the GUVs formed from DMPC and DPPC in artificial cytosol. The temperature of the electroformation was set to 45°C for DMPC and 55°C for DPPC, respectively. There was no significant GUVs formation under 5 V/10 Hz and 5 v/100 Hz AC fields for DMPC phospholipid carry artificial cytosol. The phospholipid membrane exists on the ITO glasses by adsorption mainly, which may be due to the agarose-sucrose mixed sol affecting the lipid bilayer to separate and bend. (Fig. S1a and b). For DPPC phospholipids, 5 V/10 Hz and 5 v/100 Hz AC fields can form GUVs successfully. But, the vesicle size was uneven, and there were many double vesicles in solution, as shown in Fig. S1c. One possible explanation might be the high concentration of counterions near the electrode surface and formation of an electric double layer.[30] The DPPC vesicle production is low at amplitude of 5 V and frequency of 100 Hz (Fig. S1d). GUVs formation may be suppressed if the frequency was too high.[31] The histogram of size distribution showed the heterogeneity of DPPC GUVs under 5 V/10 Hz and 5 V/100 Hz. In conclusion, neither DMPC nor DPPC is suitable for the construction of artificial cell models containing artificial cytosol.

Subsequently, DSPC with higher transition temperature was selected to prepare GUVs in artificial cytosol under AC field with amplitude of 5 V and frequency of 10 Hz at 65°C. The DSPC vesicles were uniform in size and complete in structure as shown in Fig. S2a. The size of DSPC GUVs distribution range is narrow. The proportion of GUVs in the size range about 20 μm was 58.48±1.25% (Fig. S2b). But, the *T*_m_ of DSPC is very high, the lipid membrane was inverted hexagonal II (HII) phase at 37°C, so it is necessary to add cholesterol to improve the fluidity of DSPC phospholipid membrane. A common technique used in the study of the thermodynamics of phospholipid bilayers is differential scanning calorimetry (DSC).[32–34] DSC was used to measure the DSPC, DSPC/10 mol%Chol, DSPC/15 mol%Chol, DSPC/20 mol%Chol, DSPC/30 mol%Chol GUVs including agarose-sucrose mixed sol, as shown in Fig. 2 respectively. The *T*_m_ of pure DSPC vesicles including sol was 54.8°C (Fig. 2a). With the increase of cholesterol, the *T*_m_ decreased gradually (Fig. 2b, c, d). But, when DSPC was doped with 30 mol% cholesterol, no peak appeared in the measurement range (Fig. 2e). The *T*_m_ of DSPC/20 mol%Chol dropped to 35°C, the phospholipid component could exist in dynamic flow at human body temperature of 37 °C. Therefore, DSPC/20 mol%Chol was selected as an artificial cell membrane component in agarose-sucrose@GUVs system.

**Fig. 2.**
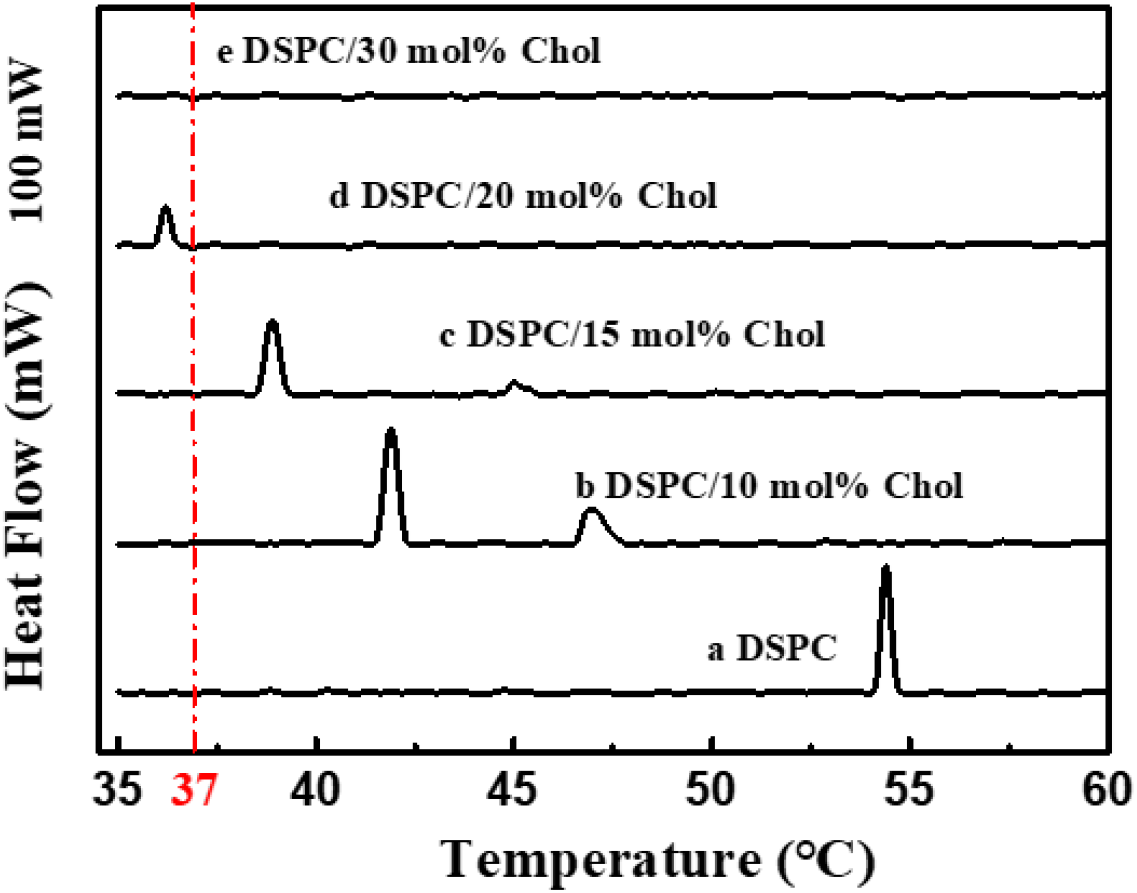
Effects of cholesterol on the *T*_m_ of DSPC.

Methods of GUVs formation include gentle hydration[35], electroformation[36, 37], solid hydration[38], and emulsion-based methods[39], etc. Among those methods, electroformation is commonly used because it generates GUVs with high yield and superior quality. There are mainly four extrinsic parameters influencing the formation of GUVs, i.e., amplitude of the field, frequency of the field. In the following, the influences from the amplitude and frequency of the AC electric field on the GUV formation were studied when the temperature was kept at 50°C. Fig. 3 (a, b, c) was the fluorescence images of different frequency and amplitude of DSPC/20 mol%Chol. The GUVs of DSPC/20 mol% Chol were formed under AC field with 5 V amplitude and 100 Hz, but the size of vesicles was small and uneven, as shown in Fig. 3a. The diameters of GUV (n=200) were assessed by analyzing the events in microscope images. The proportion of GUVs in the size range about 10 μm was 30.58±1.05% (Fig. 3d). At the same time, various other sizes can be observed in this solution. Therefore, GUVs of DSPC/20 mol% Chol were formed under AC field with 5 V amplitude and 100 Hz were not conducive to observe the subsequent experimental phenomena under the microscope. The sizes of GUV of DSPC/20 mol% Chol under AC field with 5 V amplitude and 300 Hz were relatively uniform (Fig. 3b). The size distribution range is narrow, as shown in Fig. 3e. The counts of GUVs in the size range about 15 μm was 65.08±2.63%, which was more than half. Other sizes of GUV account for a smaller proportion. The GUVs of DSPC/20 mol% Chol under AC field with 5 V amplitude and 300 Hz were suitable for the construction of artificial cell model containing cytosol. Double vesicles appeared easily of DSPC/20 mol% Chol under AC field with 5 V amplitude and 500 Hz (Fig. 3c). The size distribution of GUVs was wide as shown in Fig. 3f. In terms of both size and morphology, GUVs under this condition are not suitable for cell model to carry out subsequent experiments.

**Fig. 3.**
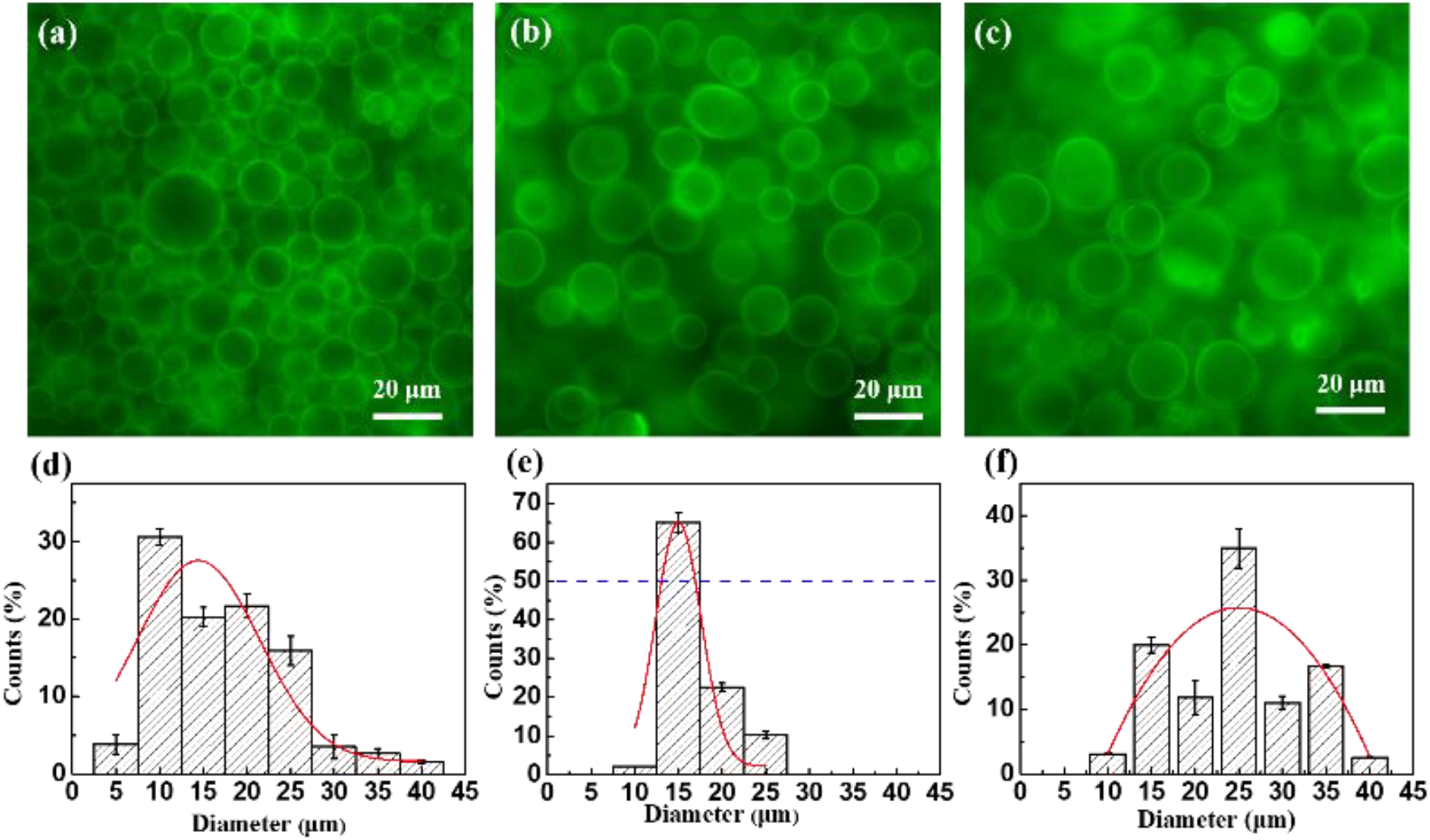
The fluorescence images of DSPC/20 mol% Chol GUVs formed on ITO electrode for 4 h in artificial cytosol under an AC field with (a) 5 V, 100 Hz, (b) 5 V, 300 Hz, (c) 5 V, 500 Hz with (d-f) their corresponding size distributions. The lipid bilayer was labelled with 1 mol% NBD-PE (green fluorescence).

### 3.3 Medical application of the agarose-sucrose@LUVs

The larger size of GUV is favorable for the observation of vesicle morphology and experimental phenomena under microscope. Rhodamine was added into water as the marker to study the release profile of DSPC/20 mol% Chol GUVs at 37°C, as shown in Fig. 4a. Rhodamine molecules in the water phase inside the vesicle can be spread to the outside of the vesicle effortlessly. When the agarose-sucrose gel was present inside the vesicle, the rate of rhodamine molecules diffusing out of the vesicle was significantly reduced. Thus, the presence of the agarose-sucrose gel facilitates the controlled-release of molecule inside the vesicle. The existence of the above conclusions was confirmed by microscope observation. Fig. 4b showed the fluorescence images of DSPC/20 mol% Chol GUVs containing rhodamine aqueous solution at pH=5. The rhodamine molecules in GUVs were obviously released completely after the GUVs was placed in this environment for 9 h. Meanwhile, Fig. 4c showed the fluorescence images of DSPC/20 mol% Chol GUVs containing rhodamine sol solution at pH=5. The fluorescence intensity in GUVs decreased slightly after 9 h, which indicated that liposome containing agarose-sucrose is suitable as a carrier to control drug release. However, GUV is too large to be used as a drug carrier. The LUV size is the best choice for the preparation of drug carriers.

**Fig. 4.**
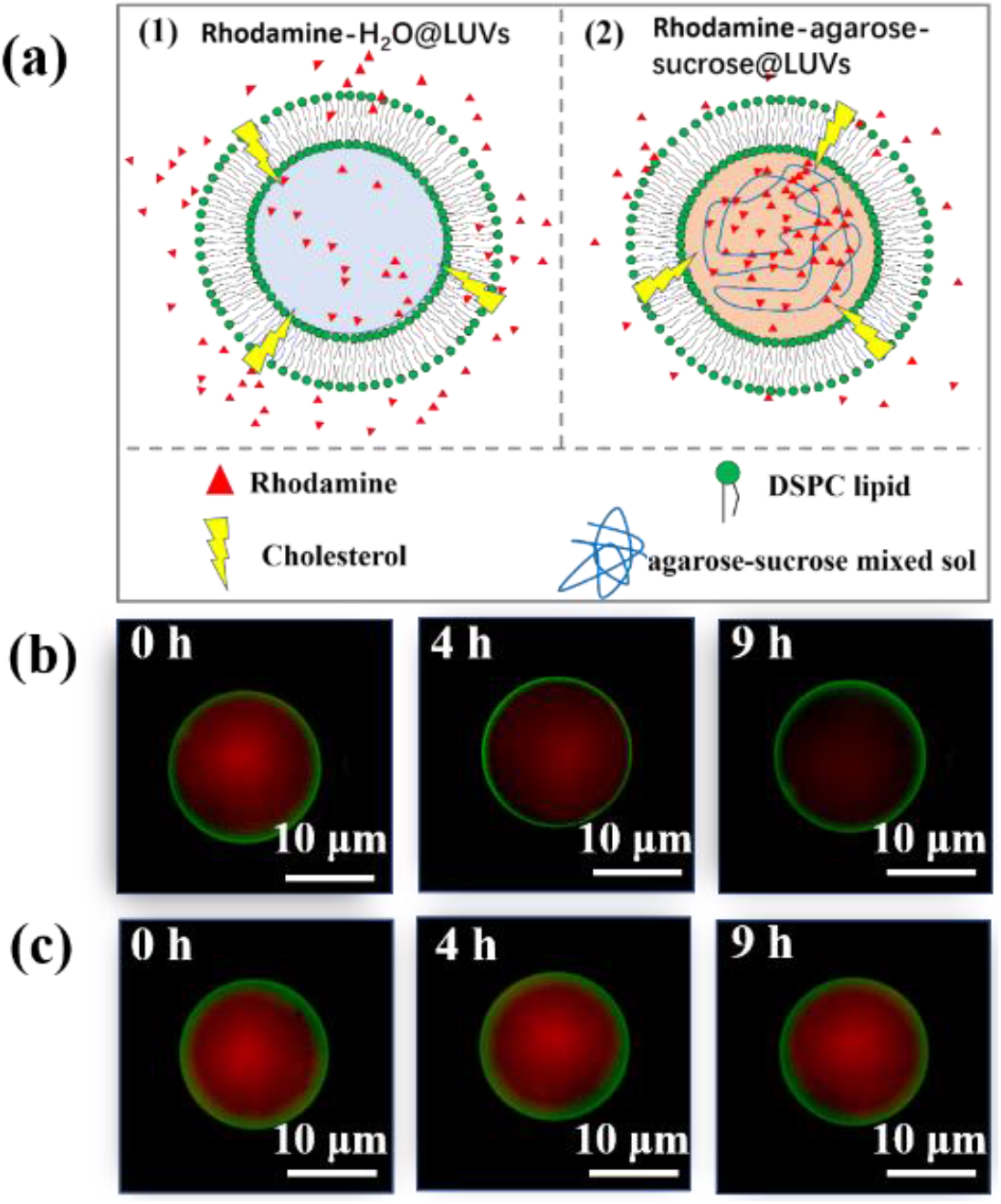
The schematic illustration of (a) water-GUVs and agarose-sucrose-GUV release rhodamine at 37°C. (b) Fluorescence images of DSPC/20 mol% Chol GUVs containing rhodamine aqueous solution and (C)DSPC/20 mol% Chol GUVs containing rhodamine agarose-sucrose gel solution at pH=5, respectively. The lipid bilayer was labelled with 1 mol% NBD-PE (green fluorescence).

Control delivery of drugs in the vector is more effective than rapid delivery of drugs directly.[40] The release profile of DOX from agarose-sucrose@LUVs exhibited a unique character compared with that from H_2_O@LUVs at lower pH solution (pH = 5). Unlike the exponential release tendency (Fig. 5a) of DOX from H_2_O@LUVs[41], the DOX release profile from agarose-sucrose@LUVs presents a persistent drug release process for a long time as shown in Fig. 5b. The release percentage of DOX-H_2_O@LUV reached a plateau at 13 h was 91.38% (Fig. 5a). At the same time, the release stage of DOX-agarose-sucrose@LUV finishes around 22 h and the total release percentage is 92.85% (Fig. 5b). They reach the similar terrace due to the same initial DOX concentration which is the predominant driven force of simple diffusion.[42] Due to the intriguing mechanism of agarose-sucrose@LUV, the release time of agarose-sucrose@LUV was prolonged 9 h more than that of H_2_O@LUVs. The existence of agarose-sucrose release behavior indicates viscous effect of colloid, which can slow down the drug release to obtain the longer drug effect in the body.

**Fig. 5.**
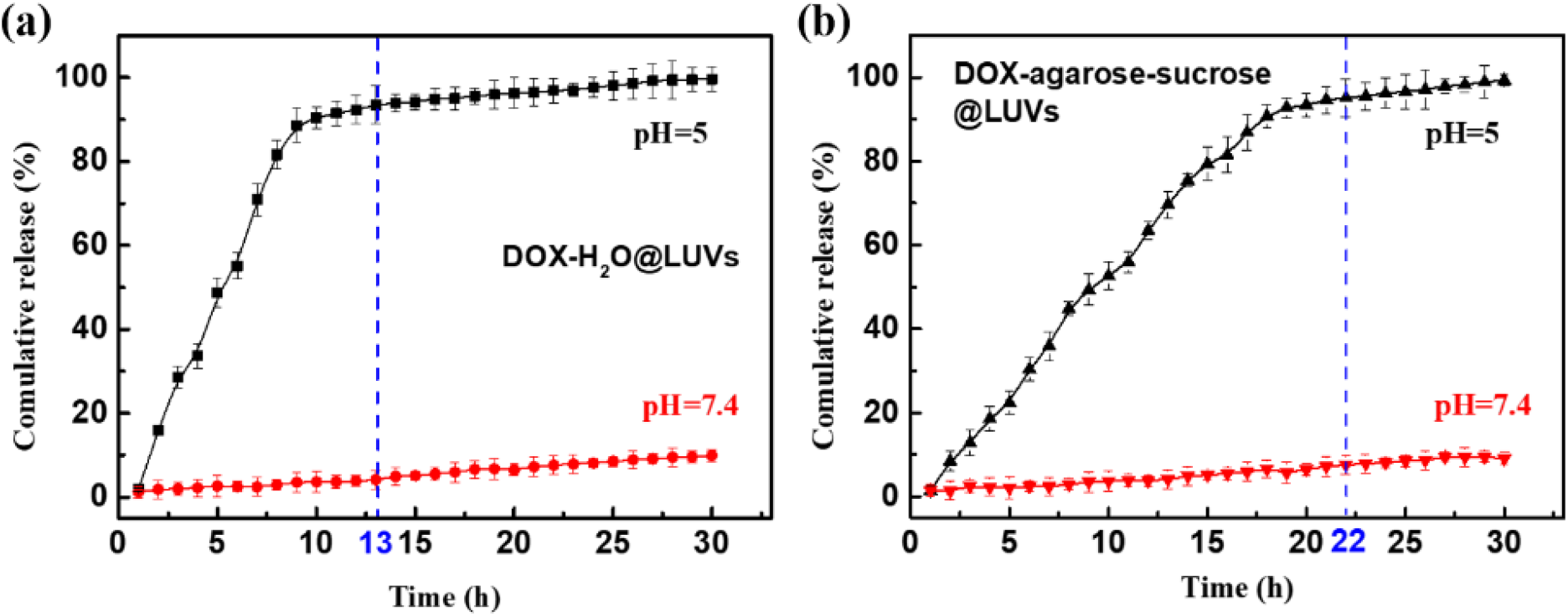
DOX release profiles of (a) DOX-H_2_O@LUVs, and (b) DOX-agarose-sucrose@LUVs. (n = 3, error bar = standard deviation).

The carriers were stored at 4 °C after producing for further experiments. In order to ensure the accuracy of the experiment, the stability of the carrier needs to be confirmed. Stabilities of various carriers were evaluated by dynamic light scattering (DLS), polymer dispersity index (PDI) and ζ-potential (SI section 3). The diameters of H_2_O@LUVs and agarose-sucrose@LUVs were about 160 nm and 100 nm, respectively, no matter whether DOX was loaded or not. The diameter and PDI of each carrier remained almost unchanged over 30 days. The values of ζ-potentials did not go down below −25 mV, which indicates the liposomes did not coagulate over a month.

The composition of agarose-sucrose@LUV was DSPC and Chol (80:20 mol%), which are biocompatible membranes. Average size of agarose-sucrose@LUV was about 125 nm (Fig. S3a) that fit in the optimum range for tumor delivery in accordance to ERP effect.[6] Inhibition of free DOX, DOX loaded LUV and DOX loaded agarose-sucrose@LUV on HeLa cells after 12, 24, 36, 48 h were summarized in Fig. 6 estimated by MTT assay[43]. As expected, the increase of DOX concentration in each group resulted in a decreased population of HeLa cells, and the cell viability was reduced with increasing incubation time. In addition, DOX@LUVs showed a higher cytotoxicity on HeLa cells than free DOX solution at each concentration.

**Fig. 6.**
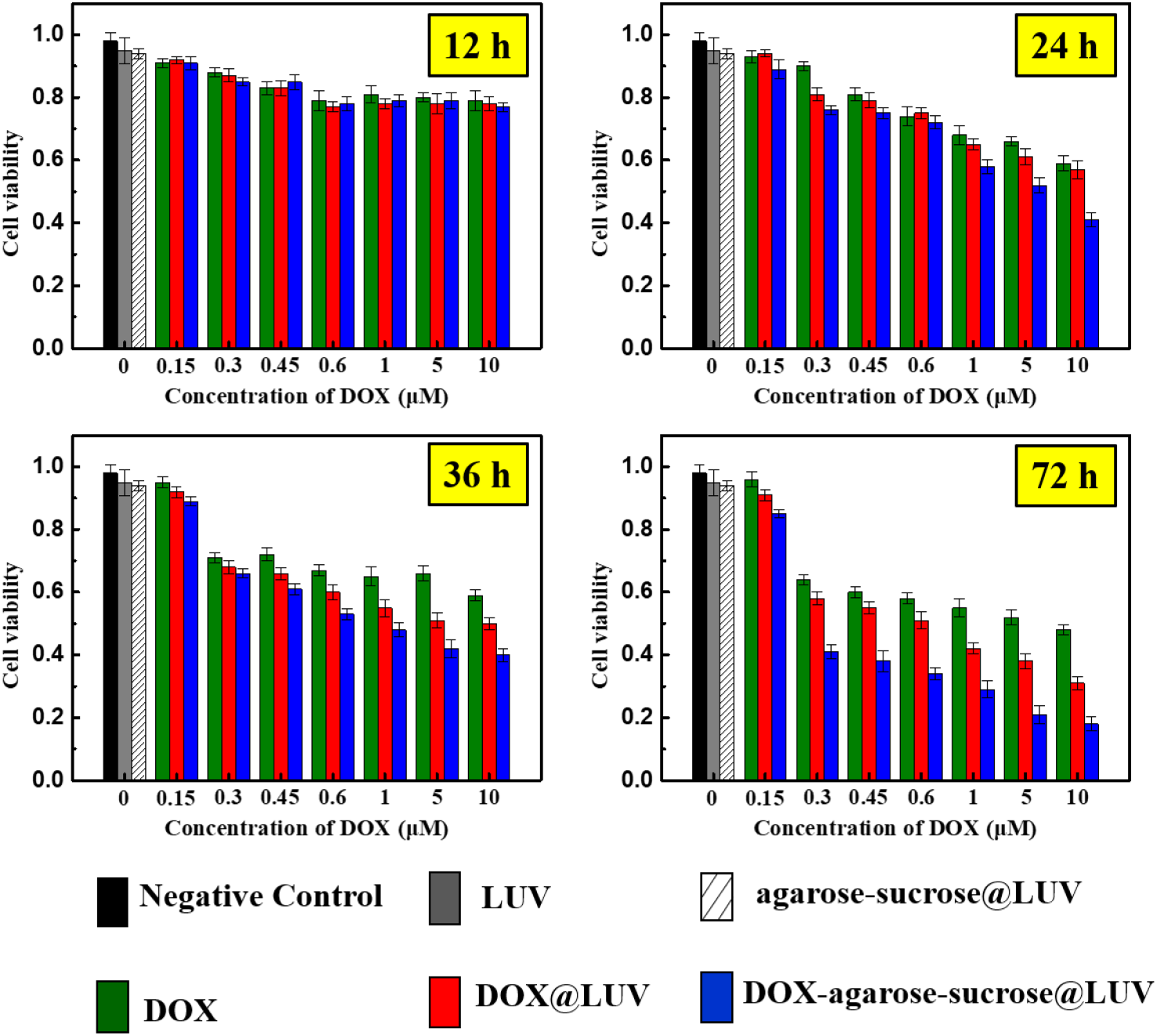
The viability of HeLa cells with treatments of control group and experimental group after 12 h, 24 h, 36 h and 48 h (n = 3, error bar = standard deviation).

Intriguingly, the performance of DOX loaded agarose-sucrose@LUVs were even better, meanwhile they showed insignificant toxicity on normal cells (SI section 4). The half maximal inhibitory concentrations (IC_50_) of DOX solution, DOX@LUV, DOX-agarose-sucrose@LUVs were 0.476 μmol L^−^ and 0.128 μmol L^−1^ and 0.015 μmol L^−1^ calculated by IBM SPSS Statistics respectively. These results demonstrated the outstanding performance of agarose-sucrose@LUVs as drug delivery system.

In order to confirm the carrier and drug distribution inside HeLa cells, NBD-PE labelled DOX-agarose-sucrose@LUVs were incubated together with HeLa cells for 12 h. Fig. 7a presented the nuclei of HeLa cells stained with DAPI. DAPI stains the nuclei compartment of both live and dead cells. Fig. 7b and c were NBD-PE and DOX distribution inside HeLa cells respectively. Fig. 7d was the merged image of Fig. 7a, b and c. From Fig. 7b, it was clearly seen that agarose-sucrose@LUVs were taken up by the tumor cells and located in their cytosol (green parts in Fig. 7b). The image in Fig. 7c indicated that the DOX are released mostly inside the cells with scarce premature release in the exterior of the HeLa cells. The LUVs can promote the uptake of DOX, and agarose-sucrose@LUVs can further escalate the uptake of DOX, as shown in Fig. 7e. The possible reason of the escalation in cytotoxicity of DOX-ATPS-LUVs is due to their slower release profile. The uptake of DOX-agarose-sucrose@LUVs was better than that of DOX@LUVs. DOX, DOX@LUVs and DOX-agarose-sucrose@LUVs of equivalent concentration 10 μM were incubated with HeLa cells for 5 h (SI section 4), separately, and determined the uptake of each group by their mean cellular fluorescence (MCF) using a flow cytometry. The MCF of each group was 267.79 ± 15.33, 384.88 ± 17.21 and 521.17 ± 25.31. These results explained the best inhibitory efficiency of agarose-sucrose@LUVs.

**Fig. 7.**
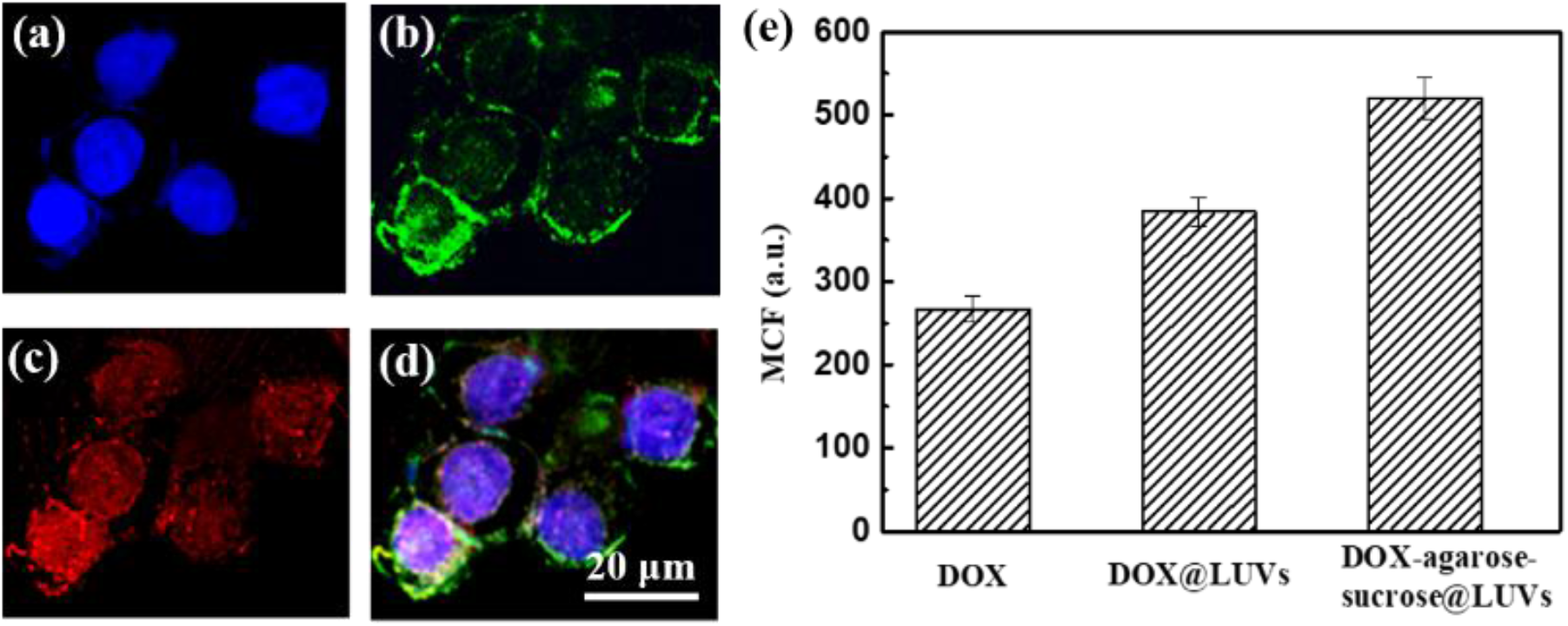
Fluorescence images of the uptake of NBD-PE labelled DOX- agarose-sucrose@LUVs by HeLa cells. (a) DAPI channel, (b) NBD-PE channel, (c) DOX channel, (d) merged image of (a), (b) and (c). (e) Mean cellular fluorescent determination of uptake of DOX, DOX@LUVs, and DOX-agarose-sucrose@LUVs by HeLa cells for 5 h.

## 4. Conclusion

We successfully presented the method of preparing artificial cytosol using agarose and sucrose. The artificial cell model was fabricated by wrapping the agarose-sucrose mixed sol in liposomes. Agarose-sucrose@LUVs exhibited a few merits as compared to free DOX and simple LUVs, such as longer sustained release time, higher uptake efficiency, meanwhile higher inhibition efficiency, which show great potential in pharmacologic therapy of cancer treatment.

## Supporting information

Supplemental manuscript

## Acknowledgements

This work was supported by the National Natural Science Foundation of China (Grant No. 22005161), the Fundamental Research Funds in Heilongjiang Provincial Universities (Grant No. 135309355, 135409212), and the Youth Training Talent’s Innovation Program of Ordinary Undergraduate Universities in Heilongjiang Province (Grant No. UNPYSCT-2020074).

---

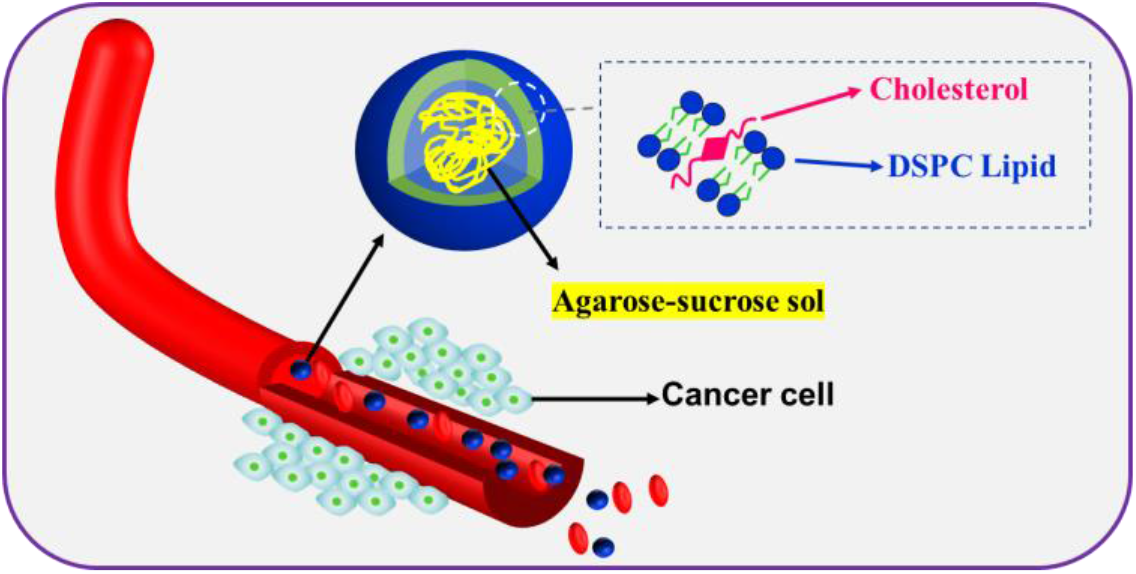
pH-sensitive liposomes encapsulating agarose-sucrose mixed sol as drug delivery carrier.

