## Supplemental manuscript for "Controllable drug release of pH-sensitive liposomes encapsulating artificial cytosol system"

### S1. Preparation of the agarose-sucrose@GUVs (DMPC) and agarose-sucrose@GUVs (DPPC)

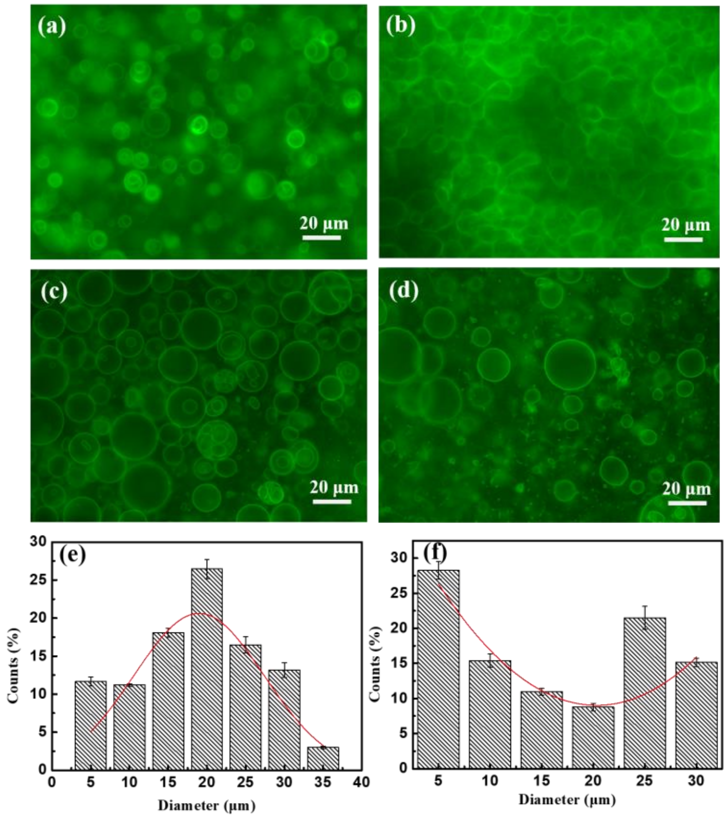


Fig. S1 Fluorescence images of GUVs formed from DMPC under 5 V amplitude and 10 Hz AC fields (a), DMPC under 5 V amplitude and 100 Hz AC fields (b), DPPC under 5 V amplitude and 10 Hz AC fields (c), and DPPC under 5 V amplitude and 100 Hz AC fields (d) in agarose-sucrose sol. The size distributions of DPPC GUVs under 5 V amplitude and 10 Hz (e), and 5 V amplitude and 100 Hz AC fields (f), respectively. The lipid bilayer was labeled with 1 mol % NBD-PE (green fluorescence).

### S2. Preparation of agarose-sucrose@DSPC-GUVs

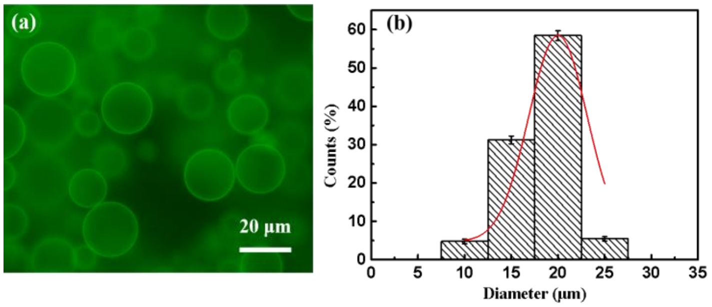


Fig. S2 Fluorescence images of GUVs formed from DSPC in agarose-sucrose sol under 5 V amplitude and 10 Hz AC fields (a) and its corresponding size distributions (b).

### S3. Estimation of stabilities

DLS and zeta-potential were taken to evaluate the stabilities of various carriers as shown in Fig. S3.

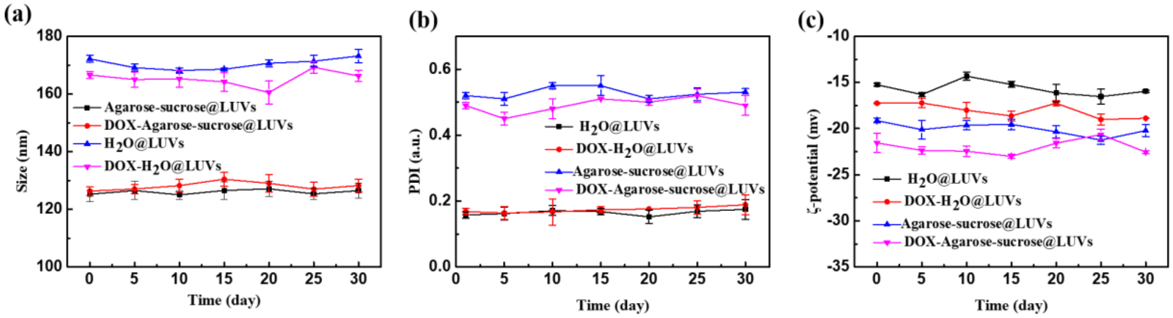


**Fig. S3.** DLS measurement of diameters (a), PDI (b) and ζ-potential (c) of each group over 30 days.

### S4. Uptake of DOX, DOX@LUVs and DOX-agarose-sucrose@LUVs

Determination of DOX, DOX@LUVs and DOX-agarose-sucrose@LUVs were estimated by flow cytometry. MCF indicates the uptake amount of DOX by HeLa cells. The uptake of each group is increasing at the beginning, and gradually dropping off after 5 h as shown in Fig. S4. The equivalent concentration of DOX was 10 μM, and the lipid concentration was 1 mg $mL^{-1}$.

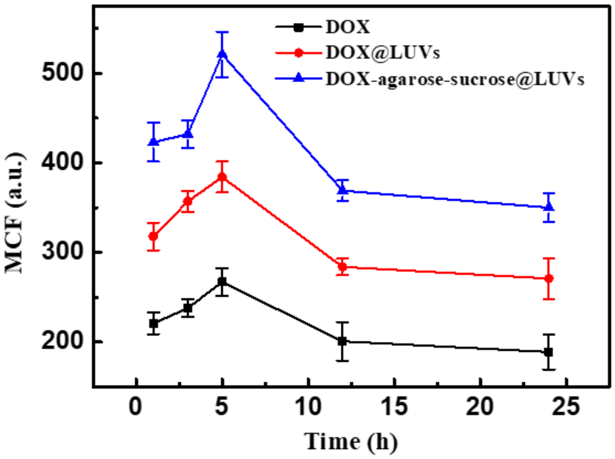


Fig. S4. MCF with respect to time of DOX, DOX@LUVs and DOX-agarose-sucrose@LUVs uptake by HeLa cells. (n=3, error bar = standard deviation)
